# Evidence of the simultaneous replications of active viruses in specimens positive for multiple respiratory viruses

**DOI:** 10.1101/2023.04.26.538472

**Authors:** Miyuki Kawase, Reiko Suwa, Satoko Sugimoto, Masatoshi Kakizaki, Yohei Kume, Mina Chishiki, Takashi Ono, Hisao Okabe, Sakurako Norito, Makoto Ujike, Mitsuaki Hosoya, Koichi Hashimoto, Kazuya Shirato

## Abstract

Genetic diagnostic assays for the detection of respiratory viruses sometimes show simultaneous multiple infections with low copy numbers. In such cases, the disease is considered caused by a single etiologic agent and others are nonspecific reactions and/or contaminations. Interferon-dependent interference is seen in dual infections of influenza and respiratory syncytial virus, which are the main causes of respiratory infections. Virus isolation is one of the solutions in detecting other active viruses present in specimens, and the air–liquid interface culture of human bronchial/tracheal epithelial cells (HBTEC-ALI) is optimal for the isolation of respiratory viruses owing to its wide range of susceptibility. In this study, we successfully confirmed the replications of various viruses from specimens with low copy numbers and passed 2–3 viruses simultaneously using HBTEC-ALI cultures, mainly including human bocavirus 1 and/or human rhinovirus.

---

Acute respiratory infections (ARIs) are the leading cause of mortality in children worldwide (1), and numerous bacteria, viruses, and fungi are associated with disease development (2-4).

The development of multiple molecular assays, such as nucleic acid amplification tests, enable the simultaneous detection of these causative agents (5-8). The coronavirus disease pandemic caused by severe acute respiratory syndrome coronavirus 2 (SARS-CoV-2), which occurred in the end of 2019, accelerated the development of assays based on real-time polymerase chain reaction (PCR) techniques (9-11). Since then, its use has become widespread among people who were not familiar with real-time PCR in virus detection. As a result, whether a high Ct in one time test indicates virus transmissibly became a complicated social problem, regardless of the difference in assays and/or amplification conditions, the time and number of diagnostic test during time course of infection (12-15).

Thus, the presence of active viruses in such multiple and/or low copy number infections should be assessed. However, in ARIs, especially in influenza (Flu) and respiratory syncytial viruses (RSV), viruses interfere with each other by inducing interferon-related genes; therefore, the virus that first caused an infection defeated the second virus (16-18). Therefore, the disease is considered caused by a single etiologic agent, and the effects of multiple infections on disease development are controversial (5, 8, 19).

Virus isolation is one of the methods of assessing the presence of active infectious agents in such suspicious situations. The air–liquid interface culture of human bronchial/tracheal cells (HBTEC-ALI) is an excellent tool for culturing various respiratory pathogens, particularly those that are hard to isolate, such as human bocavirus 1 (HBoV1) and human rubulavirus 4 [human parainfluenzavirus (HPIV4)], due to the wide range of susceptibility for respiratory viruses (20, 21). In our laboratory, virus isolations using HBTEC-ALI cultures have been performed with clinical specimens obtained from pediatric in-patients with severe ARI (SARI) (22, 23).

In this study, we aimed to report that we successfully isolated various viruses from specimens with low copy number, and we detected different viruses simultaneously, mainly including HBoV1 and/or human rhinovirus (HRV).

## Methods

### Clinical specimens

Clinical specimens were collected in acute phase from pediatric in-patients with SARIs in Fukushima, Japan, between 2018 and 2022 (22). Nasopharyngeal swabs were collected in the universal transport medium (Copan, Brescia, Italy) and stored at −80°C until use.

The study protocol was approved by the Ethics Committees of the National Institute of Infectious Diseases (Nos. 1001, 1087, and 1441) and Fukushima Medical University (No. 29006). Informed consent was obtained verbally and was noted in the medical records in each hospital. In this study, SARS-CoV-2-negative specimens were used to evaluate respiratory infections as common diseases.

### Virus isolation

Nucleic acids were extracted from specimens using QIAamp 96 Virus QIAcube HT Kit (Qiagen, Hilden, Germany), QIAamp Viral RNA Mini Kit (Qiagen), or NucleoSpin 96 Virus (Macherey-Nagel, Duren, Germany) following the manufacturer’s instruction, except that the elusion step was performed by centrifugation. Respiratory viruses were detected from specimens by real-time PCR and reverse transcription (RT)-PCR assays with LightCycler instruments (Roche, Basel, Switzerland) as previously described (22, 24). The real-time PCR value was expressed as crossing point (Cp). Primer/probe sequences are listed in Table S1. The following 17 viruses were tested: human orthopneumovirus (RSV subgroups A and B); Flu A, B, and C; human coronavirus (HCoV) (229E, OC43, NL63, and HKU1); human metapneumovirus (hMPV); HPIV (1, 2, 3, and 4); adenovirus (ADV) [2 (for A, C, D, and F) and 4 (B and E)]; HBoV1; and HRV (6, 23, 25-27). Among the above specimens, 249 specimens were positive for respiratory viruses (single infection, n = 122; multiple infections, n = 127) and were subjected to virus isolation. The air–liquid interface culture of human bronchial/tracheal epithelial cells (HBTEC-ALI) was prepared as described previously (20, 21, 28). Then, 20 µL of the specimen diluted with an equal volume of 1%FCS-DMEM containing prescribed amount of antibiotics (penicillin-streptomycin, gentamicin, and fungizone) was inoculated onto the apical surface of HBTEC-ALI cultures. Cells were incubated at 34°C overnight and then washed with 1%FCS-DMEM four times. The fourth wash solution was kept as the starting control. The MC-210 (KAC Co Ltd., Hyogo, Japan) was added to the basolateral medium throughout the incubation periods to remove mycoplasmas and every buffer change a week. After 4, 7, 11, 18, and 25 days of incubation at 34°C (18 and 25 days were depended on the condition of culture cells), cells were washed with 1%FCS-DMEM four times, and the cell-washes were stored at −80°C. Virus replications were confirmed by real-time RT-PCR assays as described above (22). During the time course and passage, when the specimen showed >3.3 Cp (≈ 1 log_10_) of increase relative to day 0 and/or 4, it was considered positive virus isolation. The copy number of each virus was determined using the calibration curve drawn by real-time RT-PCR assay with concentration-calculated synthesized control DNA or RNA containing the primer/probe targeted region. The copy numbers of the control DNA/RNA were calculated based on the molecular weight and absorbance at 260 nm of RNA that had been serially diluted (≥3 steps) and/or fluorescence using the Quantus Fluorometer (Promega, Madison, WI, USA) (n = 3).

### Sequencing analysis

For next-generation sequencing, libraries were prepared using the NEBNext Ultra II RNA Library Prep Kit for Illumina [New England Biolabs (NEB), Ipswich, MA, USA] or the NEBNext Ultra II FS DNA Library Prep Kit for Illumina (NEB), following the manufacturer’s instructions. Indexed libraries were analyzed for 2×150 cycles on a DNBSEQ-G400 sequencer at AZENTA/GENEWIZ (Chelmsford, MA, USA). Reads were trimmed and then *de novo* assembled and/or mapped to each reference sequence using the CLC Genomics Workbench (v21.0.4, Qiagen) with default settings.

### Statistical analysis

Statistical analysis was performed using SigmaPlot software (Ver. 14.5, Systat Software Inc., Palo Alto, CA, USA). The *z*-test and chi-square test were used. The *t*-test or the Mann-Whitney test were also used depending on the normality test result. The *p* value less than 0.05 was considered as significant.

## Results

### Respiratory virus isolation using HBTEC-ALI cultures

The real-time (RT-)PCR techniques sometimes detect multiple positives for various targets in the tested specimen with suspicious respiratory infections. Virus isolation is the most suitable technique in identifying the “active” virus in specimens positive for multiple agents. Furthermore, HBTEC-ALI cultures are the best cells for respiratory virus isolation owing to the wide range of susceptibility. In this study, HBTEC-ALI cultures were used, and virus isolation was performed with clinical specimens obtained from pediatric in-patients with respiratory infections (22, 23). In HBTEC-ALI culture, cells generally did not show a cytopathic effect; therefore, genetic replication detected by real-time RT-PCR was used as the indicator of successful virus isolation. The replication kinetics of virus should be monitored by repeated test during time course of infection, in initial diagnosis, only one time test result can be available, generally. Therefore, the results of real-time (RT-)PCR in initial diagnosis were applied following experiment. The 249 specimens (122 single infection and 127 multiple infections) were subjected to virus isolation, and 131 specimens (52.6%) were positive for replication of some viruses. Moreover, 70 of 122 single-infection specimens were positive (57.3%) and 61 of 127 multiple-infection specimens were positive in at least one target (48.0%). No significant differences in virus isolation were found between single- and multiple-infection specimens by *z*-test (*p* = 0.177). For 17 respiratory viruses, 450 targets were tested, and 147 of these (32.7%) showed virus replication (Table 1). The rates of successful virus replication were low in RSVA (*p* = 0.04) and higher in FluA (*p* = 0.014) and FluB (*p* = 0.002) than average. Overall, successful virus replication was related to lower real-time RT-PCR value (25.7 vs 33.0, Cp value), i.e., high amounts of inoculated virus led to positive results (4.6 vs 2.5 log_10_ copies, *p* < 0.001) (Table 1). Yamada et al., reported that 87.5 % of successful virus isolation was seen in less than 28.1 of Cp, which was corresponding to about log 3.5 copies determined by NIID-N2 set, in the isolation of SARS-CoV-2 using VeroE6/TMPRSS2 cells (29). The average (±confidence interval) copies of Cp = 28.1 in used real-time PCR assays is log 3.23±0.36. Although the correlation between real-time PCR value and viral copy number is completely dependent on the performance of each primer/probe set (the same Cp value does not always show the same copy number in different primer/probe sets), data were sorted by the Cp value (≥28.1) to evaluate successful virus replication in low-copy specimens (Table 2). In total, among 296 targets, 54 showed virus replication (18.2%). The replication rates of FluB, and HPIV1 were higher than average. In total, successful replications were also seen in specimens with lower Cp values (31.8 vs 34.9, *p* < 0.001) and the higher copy number (2.8 vs 1.9, log_10_ copies, *p* < 0.008). The performance of each real-time (RT-)PCR assays are different and same Cp value does not always show same copy number; therefore, the results have been converted to the copy number enable to compare altogether. In addition, the sensitivity of virus replication on HBTEC-ALI culture are different depending on the virus species. However, these results indicate the tendency that the higher the number of viral copies, the more successful is the virus replication. Although approximately several hundred of copies are required to lead to possible respiratory virus isolation in HBTEC-ALI cultures, intact virion (active virus) surely exists even in low copies specimens,

**Table 1.** Results of virus isolation

|  | Target<sup>a</sup> | RSVA | RSVB | FluA | FluB | HCoV<br>229E | HCoV<br>OC43 | HCoV<br>NL63 | HCoV<br>HKU1 | hMPV | PIV1 | PIV2 | PIV3 | PIV4 | FluC | ADVC | ADVB3 | HBoV | HRV | Total |
| --- | --- | --- | --- | --- | --- | --- | --- | --- | --- | --- | --- | --- | --- | --- | --- | --- | --- | --- | --- | --- |
|  | Total test | 39 | 25 | 13 | 6 | 10 | 24 | 18 | 13 | 48 | 10 | 7 | 26 | 35 | 6 | 44 | 16 | 53 | 57 | 450 |
|  | Replicated | 6 | 4 | 9 | 6 | 2 | 12 | 8 | 3 | 14 | 6 | 4 | 11 | 10 | 1 | 8 | 5 | 18 | 20 | 147 |
|  | Success rate<br>(%) | 15.4 | 16.0 | 69.2 | 100.0 | 20.0 | 50.0 | 44.4 | 23.1 | 29.2 | 60.0 | 57.1 | 42.3 | 28.6 | 16.7 | 18.2 | 31.3 | 34.0 | 35.1 | 32.7 |
| Average Cp | Replicated | 22.4 | 20.7 | 24.2 | 29.5 | 24.2 | 23.7 | 26.0 | 27.6 | 26.9 | 31.2 | 25.6 | 27.2 | 31.1 | 21.1 | 24.7 | 23.0 | 22.9 | 31.2 | 25.7 |
|  | Non-replicated | 25.7 | 28.7 | 30.2 | N.A<sup>b</sup> | 36.4 | 32.1 | 30.6 | 35.6 | 31.2 | 29.4 | 37.1 | 36.7 | 35.3 | 31.3 | 35.5 | 37.1 | 34.8 | 33.8 | 33.0 |
| Inoculated<br>copies (log) | Replicated | 5.6 | 6.2 | 4.3 | 3.2 | 5.4 | 4.3 | 4.5 | 3.6 | 5.2 | 2.2 | 4.6 | 5.6 | 2.8 | 6.1 | 4.6 | 5.6 | 5.2 | 4.2 | 4.6 |
|  | (Range) | 3.6-6.8 | 5.6-6.5 | 2.0-6.1 | 2.5-3.6 | 4.3-6.5 | 2.4-6.8 | 2.9-6.0 | 1.3-6.5 | 4.3-6.5 | 0.9-3.6 | 2.7-6.0 | 4.3-7.3 | 0.8-4.5 | N.A | 1.1-6.7 | 1.5-7.8 | 2.3-7.7 | 2.4-6.6 |  |
|  | Non-replicated | 4.6 | 4.0 | 2.5 | N.A | 1.4 | 1.7 | 3.2 | 1.1 | 3.9 | 2.7 | 1.2 | 2.8 | 1.7 | 3.1 | 1.2 | 1.6 | 1.5 | 3.5 | 2.5 |
|  | (Range) | 1.3-6.9 | 1.8-7.3 | 2.0-3.7 | N.A | 0.6-2.9 | 0.4-5.0 | 1.2-5.5 | 0.2-2.6 | 1.4-6.7 | 1.0-4.7 | 0.8-1.8 | 1.8-5.3 | 0.4-4.3 | 1.5-5.3 | -0.3-4.2 | 0.8-2.8 | 0.0-4.1 | 2.0-7.6 |  |
|  | *P* value<sup>c</sup> | 0.33 | <0.001 | 0.042 | N.A | <0.151 | <0.001 | 0.06 | 0.245 | <0.001 | 0.595 | 0.018 | <0.001 | 0.019 | N.A | <0.001 | 0.015 | <0.001 | 0.03 | <0.001 |

a: ADV2 set detects A, C, D, and F and ADV4 set detects B and E.
b: N/A, not applicable
c: Statistical differences in copy number between replicated and non-replicated specimens were analyzed by *t* test or Mann-Whitney test on SigmaPlot depending on the result of normalization test.

**Table 2.** Results of virus isolation (Cp ≥ 28.1)

|  | Target<sup>a</sup> | RSVA | RSVB | FluA | FluB | HCoV<br>229E | HCoV<br>OC43 | HCoV<br>NL63 | HCoV<br>HKU1 | hMPV | PIV1 | PIV2 | PIV3 | PIV4 | FluC | ADV2 | ADV4 | HBoV | Rhino | Total |
| --- | --- | --- | --- | --- | --- | --- | --- | --- | --- | --- | --- | --- | --- | --- | --- | --- | --- | --- | --- | --- |
|  | Total test | 11 | 11 | 5 | 4 | 8 | 11 | 10 | 12 | 27 | 7 | 5 | 20 | 31 | 4 | 37 | 12 | 32 | 49 | 296 |
|  | Replicated | 0 | 0 | 2 | 4 | 0 | 1 | 3 | 2 | 4 | 5 | 2 | 5 | 7 | 0 | 2 | 1 | 2 | 14 | 54 |
|  | Success rate<br>(%) | 0.0 | 0.0 | 40.0 | 100.0 | 0.0 | 9.1 | 30.0 | 16.7 | 14.8 | 71.4 | 40.0 | 25.0 | 22.6 | 0.0 | 5.4 | 8.3 | 6.3 | 28.6 | 18.2 |
| Average | Replicated | N.A.<sup>b</sup> | N.A. | 31.5 | 30.3 | N.A. | 29.9 | 31.0 | 32.3 | 30.3 | 32.2 | 30.1 | 30.3 | 32.9 | N.A. | 32.5 | 37.5 | 30.9 | 34.1 | 31.8 |
| Cp | Non-replicated | 33.9 | 34.2 | 31.5 | N.A. | 36.4 | 33.8 | 33.5 | 35.6 | 33.8 | 33.9 | 37.1 | 36.7 | 35.6 | 33.1 | 35.8 | 37.1 | 36.1 | 34.6 | 34.9 |
| Inoculated<br>copies<br>(log) | Replicated | N.A. | N.A. | 2.1 | 3.0 | N.A. | 2.4 | 3.1 | 2.2 | 4.2 | 1.9 | 3.3 | 4.7 | 2.3 | N.A. | 2.1 | 1.5 | 2.7 | 3.5 | 2.8 |
|  | Non-replicated | 2.1 | 2.6 | 2.1 | N.A. | 1.4 | 1.2 | 2.4 | 1.1 | 3.2 | 1.4 | 1.2 | 2.8 | 1.6 | 2.6 | 1.1 | 1.6 | 1.1 | 3.3 | 1.9 |

a: ADV2 set detects A, C, D, and F and ADV4 set detects B and E.
b: N/A, not applicable

### Simultaneous virus replication in HBTEC-ALI cultures

In the virus isolation, 16 specimens showed simultaneous replications of 2–3 viruses in the first inoculation in the same HBTEC-ALI culture (Table S2). Eleven of them were in combination with HBoV1 or HRV, and three were in combination with ADV2. Combinations of envelope viruses were seen in two (HCoV229E/HMPV and HPIV1/HPIV4). Eleven of the co-replicated viruses could be simultaneously passed once (Table 3 and 4). In the H739 specimen, ADVC, ADVB3, and HRV were replicated after 7 days in the first isolation (P0) despite a high Cp value in ADVC and HRV, and both ADVC and ADVB3 could be passed one time (P1). In the O898 specimen, ADVC and ADVB3 were replicated simultaneously in both P0 and P1 cultures. In the O876 specimen, HPIV1 and HPIV4 were replicated and passed in both P0 and P1 cultures despite high Cp values in the specimen. In the OR7 specimen, HCoVOC43, HRV, and HBoV1 were replicated in the P0 culture, and HCoVOC43 and HRV could be passed in the P1 culture. Simultaneous replications and passage of HBoV1 and HRV were most observed in specimens OR291, OR321, and OR331. The following four specimens could be simultaneously passed two times (Table 4). In the H257 specimen, HCoVNL63 and HBoV1 were replicated throw the passage. In the H260 specimen, although HBoV1 was replicated in the P1 culture, HPIV3 and HRV were replicated during two passages. In the OR32 specimen, ADVC and HBoV1 were replicated and passed till the P2 culture. In the OR65 specimen, HBoV1 and HRV were co-replicated in all passage. In these passage-successful specimens, the nearly complete genomic sequencers of these replicated viruses could be decoded and were registered in the GenBank database (Table 5), proofing the presence of active viruses simultaneously in the same cell cultures. These can be strong evidence that the specimens showing positivity for multiple agents in real-time PCR tests surely contain infectious viruses.

**Table 3.**
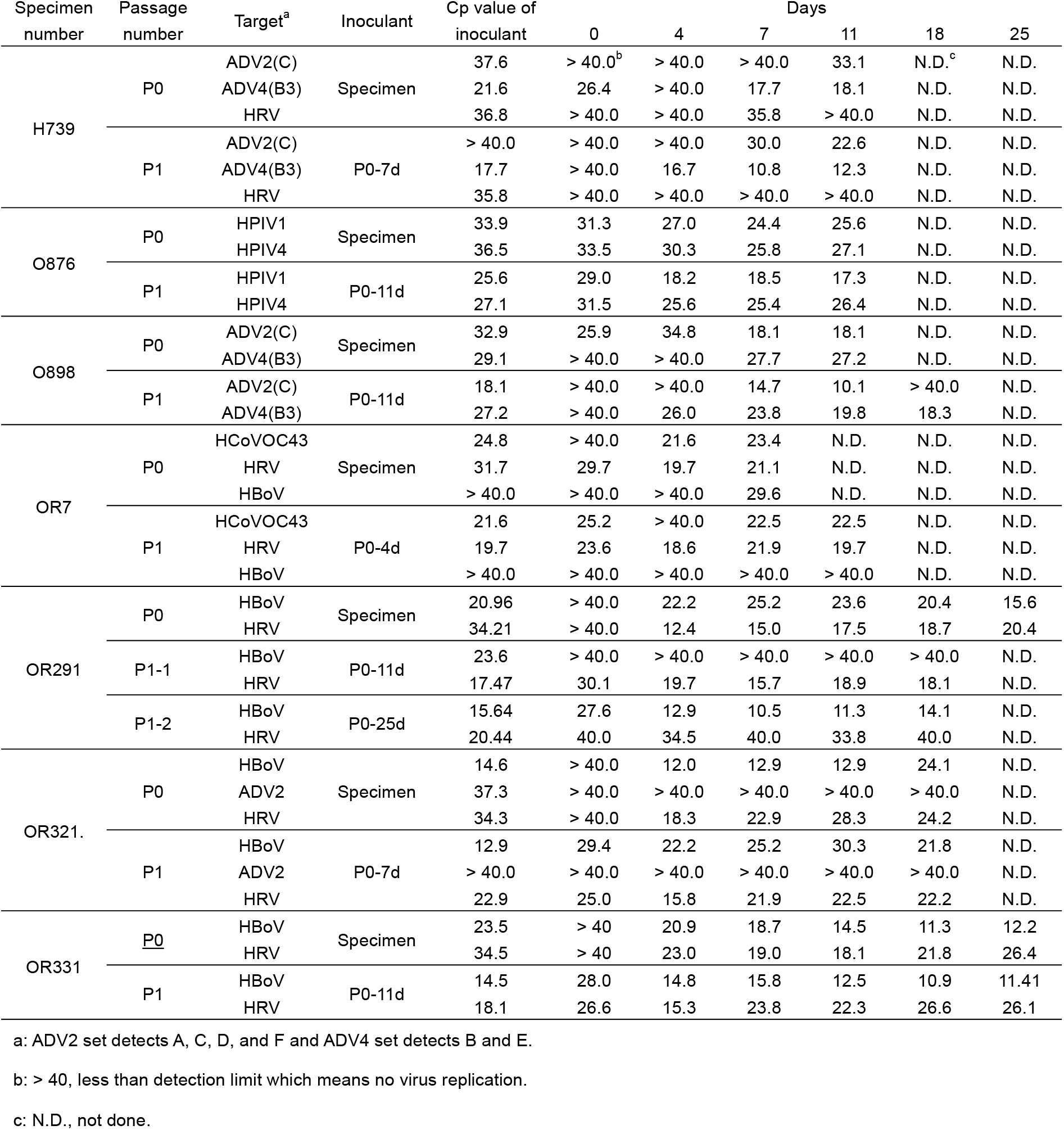
Simultaneous replication of multi-positive specimens.

**Table 4.** Serial passage of multi-positive specimens.

| Specimen number | Passage number | Target<sup>a</sup> | Inoculant | Cp value of inoculant | Days |  |  |  |  |
| --- | --- | --- | --- | --- | --- | --- | --- | --- | --- |
|  |  |  |  |  | 0 | 4 | 7 | 11 | 18 |
| H257 | P0 | RSV | Specimen | 30.7 | > 40.0<sup>b</sup> | > 40.0 | > 40.0 | > 40.0 | N.D.<sup>c</sup> |
|  |  | HCoVNL63 |  | 31.0 | 38.2 | 16.8 | 17.7 | 15.6 | N.D. |
|  |  | HBoV |  | 17.2 | 22.5 | 20.7 | 26.0 | 26.7 | N.D. |
|  | P1 | RSV | P0-7d | 40.0 | > 40.0 | > 40.0 | > 40.0 | > 40.0 | > 40.0 |
|  |  | HCoVNL63 |  | 17.7 | 28.8 | 19.0 | 17.2 | 17.9 | 15.4 |
|  |  | HBoV |  | 26.0 | 39.5 | 36.7 | 31.4 | 38.9 | > 40.0 |
|  | P2 | RSV | P1-7d | 40.0 | > 40.0 | > 40.0 | > 40.0 | > 40.0 | > 40.0 |
|  |  | HCoVNL63 |  | 17.2 | 40.0 | 19.6 | 17.8 | 23.0 | 22.4 |
|  |  | HBoV |  | 31.4 | 37.9 | 17.3 | 14.6 | 13.7 | 12.2 |
| H260 | P0 | HPIV3 | Specimen | 29.9 | 35.3 | 37.5 | 19.0 | 19.2 | N.D. |
|  |  | HBoV |  | 33.0 | 37.9 | > 40.0 | > 40.0 | > 40.0 | N.D. |
|  |  | HRV |  | 38.1 | 38.9 | 27.9 | 21.7 | 24.0 | N.D. |
|  | P1 | HPIV3 | P0-4d | 19.0 | 29.1 | 22.3 | 22.1 | 23.3 | N.D. |
|  |  | HBoV |  | > 40.0 | > 40.0 | > 40.0 | > 40.0 | 29.6 | N.D. |
|  |  | HRV |  | 21.7 | 27.6 | 24.7 | 24.1 | 23.6 | N.D. |
|  | P2 | HPIV3 | P1-11d | 23.3 | 31.6 | 25.5 | 28.4 | N.D. | N.D. |
|  |  | HBoV |  | 29.6 | 31.8 | > 40.0 | > 40.0 | N.D. | N.D. |
|  |  | HRV |  | 23.6 | 38.5 | 33.1 | > 40.0 | N.D. | N.D. |
| OR32 | P0 | ADV2 (C) | Specimen | 21.8 | 28.9 | 17.4 | 24.3 | N.D. | N.D. |
|  |  | HBoV |  | 27.7 | > 40.0 | 37.8 | > 40.0 | N.D. | N.D. |
|  |  | HRV |  | 36.1 | > 40.0 | > 40.0 | > 40.0 | N.D. | N.D. |
|  | P1 | ADV2 (C) | P0-4d | 17.4 | 27.8 | 38.3 | 21.6 | 19.1 | N.D. |
|  |  | HBoV |  | 37.8 | > 40.0 | 23.2 | > 40.0 | > 40.0 | N.D. |
|  |  | HRV |  | > 40.0 | > 40.0 | 38.5 | > 40.0 | > 40.0 | N.D. |
|  | P2 | ADV2 (C) | P1-4d | 38.3 | > 40.0 | > 40.0 | 14.0 | 11.0 | > 40.0 |
|  |  | HBoV |  | 23.2 | > 40.0 | > 40.0 | > 40.0 | 25.3 | 16.2 |
|  |  | HRV |  | 38.5 | > 40.0 | > 40.0 | > 40.0 | 38.1 | > 40.0 |
| OR65 | P0 | HBoV | Specimen | 23.4 | 30.0 | 17.7 | 12.7 | N.D. | N.D. |
|  |  | HRV |  | 29.6 | 28.5 | 20.6 | 21.5 | N.D. | N.D. |
|  | P1 | HBoV | P0-7d | 12.7 | 22.1 | > 40.0 | 14.7 | 16.4 | N.D. |
|  |  | HRV |  | 21.5 | 30.6 | > 40.0 | 28.8 | 29.3 | N.D. |
|  | P2 | HBoV | P1-7d | 10.1 | 29.1 | 17.1 | 10.1 | 10.8 | N.D. |
|  |  | HRV |  | 24.2 | 33.1 | 23.2 | 24.2 | 30.6 | N.D. |

a: ADV2 set detects A, C, D, and F and ADV4 set detects B and E.
b: > 40, less than detection limit which means no virus replication.
c: N.D., not done.

**Table 5.** GenBank accession numbers for nearly complete genomic sequences of simultaneous passaged viruses.

| Specimen number | Isolate name | GenBank accession number |
| --- | --- | --- |
| H739 | ADV2_Fukushima_H739_2019 | LC756665 |
|  | ADVB3_Fukushima_H739_2019 | LC703523 |
|  | HRV_A15_Fukushima_H739_2019 | LC756672 |
| O876 | PIV1_Fukushima_O876_2019 | LC720863 |
|  | PIV4b_Fukushima_O876_2019 | LC720882 |
| O898 | ADV2_Fukushima_O898_2019 | LC756666 |
|  | ADVB3_Fukushima_O898_2019 | LC757030 |
| OR7 | HCoVOC43_Fukushima_OR7_2020 | LC720427 |
|  | HRV_A40_Fukushima_OR7_2020 | LC720413 |
|  | HBoV_Fukushima_OR7_2020 | LC720423 |
| OR291 | HBoV_Fukushima_OR291_2021 | LC720417 |
|  | HRV_A80_Fukushima_OR291_2021 | LC720411 |
| OR321 | HBoV_Fukushima_OR321_2021 | LC720419 |
|  | HRV_C6_Fukushima_C6_2021 | LC720414 |
| OR331 | HBoV_Fukushima_OR331_2021 | LC720422 |
|  | HRV_A80_Fukushima_OR331_2021 | LC720412 |
| H257 | HCoVNL63_Fukushima_H257_2018 | LC687394 |
|  | HBoV_Fukushima_H257_2018 | LC720416 |
| H260 | PIV3_Fukushima_H260_2018 | LC720871 |
|  | HRV_A78_Fukushima_H260_2018 | LC699414 |
| OR32 | HAdVC_Fukushima_OR32_2020 | LC720426 |
|  | HBoV_Fukushima_OR32_2020 | LC720421 |
| OR65 | HBoV_Fukushima_OR65_2020 | LC651178 |
|  | HRV_A80_Fukushima_OR65_2020 | LC699423 |

## Discussion

In this study, we show evidence on the presence of an active virus in specimens with low copies and/or positive for multiple respiratory infections. The results clearly show successful virus replication in HBTEC-ALI cultures inoculated with specimens with Cp >28.1 and simultaneous replications and serial passages of 2–3 viruses, mainly including HBoV1 and/or human rhinovirus.

During the COVID-19 pandemic, real-time RT-PCR-based diagnostic assays have been used to detect SARS-CoV-2; however, the correlation between positive signals in real-time RT-PCR and the possibility of virus spread from patients became a problem because PCR-negative results had been required for discharge in the early phase of pandemic (30-32). For SARS-CoV-2, the assays that targeted structural proteins such as nucleocapsid gene were well-used (9, 11); however, these assays also detect subgenomic mRNAs of coronaviruses, causing confusion on the presence of intact virions. In a previous report, the relation between real-time RT-PCR and successful virus isolation was assessed using the ORF1a assay, which could detect mainly genomic RNA, and several dozens of copies of genomic RNA were required for possible virus isolation (33). In the present study, various real-time RT-PCR assays were used, and comparing their results using the value of real-time PCR (such as Ct and Cp) was difficult. However, it enables conversion of the results to copy numbers, and the presence of approximately several hundred copies of viruses possibly leads to replications in HBTEC-ALI cultures for all respiratory viruses tested in this study. In suspicious specimens, the copy number, not the RT-PCR value, should be used as a guide for possible infectivity. Note that the frozen specimens were used in virus isolation in this study due to the convenience of transportation, and it might affect the result of virus replication like RSV.

Multiple respiratory viruses infect respiratory tissues one after another and induce reciprocal interference in one another, i.e., Flu, RSV, HMPV, and HRV interfere with other viruses by the signaling of intracellular RNA sensors or inducing interferon-related genes (16, 17). This is also a major theory in hepatitis, i.e., hepatitis C virus infection inhibits the infections of other hepatitis viruses (34, 35). Therefore, we tended to think that the first virus suppresses the second virus, and the infectious diseases were mainly caused by the first virus. Certainly, only one virus could be replicated in most specimens with multiple infections (73.8%, 45 of 61, Table S2); however, simultaneous virus replications were observed, although mainly with HBoV1, HRV, and/or ADV infections. Several report described persistent infection of HBoV1, HRV, and ADV might cause high replication rate of these viruses (36-38). HBoV1 and HRV cause severe respiratory tract infections with simultaneous infections caused by other infectious agents (39-42). By contrast, some reports described that co-infections with these viruses do not affect the disease severity (43, 44). Thus, the relationship between the synergistic replication of infected dual viruses and disease severity must be investigated. In addition, the mechanism of acceptable dual virus replication is unclear. Although many reports describe one virus infection disturbs other virus infections (16-18), there is no report for viral symbiotic. Therefore, it is necessary to elucidate the acceptable combination of respiratory virus species for dual virus replication and the mechanisms. Furthermore, in this study, although the co-cultivative cases are minority, it is due to the limit of the sensitivity of virus isolation system. Improvement of HBTEC-ALI culturing system might increase the number of dual replication cases.

## Acknowledgments

We thank Masatoki Sato, Department of Pediatrics, School of Medicine, Fukushima Medical University; Hiroko Sakuma, Hoshi General Hospital; and Shigeo Suzuki, Ohara General Hospital, for the collection of clinical specimens. This work was supported by Grants-in-Aid (22fk0108119j0603 and 22kf0108117j0103) from the Japan Agency for Medical Research and Development and by a Grant-in-Aid from the Japan Society for the Promotion of Science (C:20K06441).

## Data availability

The nearly complete genome sequences of isolates have been deposited in GenBank under the following accession numbers (Table 5): <u>LC756665</u>, <u>LC703523</u>, <u>LC756672</u>, <u>LC720863</u>, <u>LC720882</u>, <u>LC756666</u>, <u>LC757030</u>, <u>LC720427</u>, <u>LC720413</u>, <u>LC720423</u>, <u>LC720417</u>, <u>LC720411</u>, <u>LC720419</u>, <u>LC720414</u>, <u>LC720422</u>, <u>LC720412</u>, <u>LC687394</u>, <u>LC720416</u>, <u>LC720871</u>, <u>LC699414</u>, <u>LC720426</u>, <u>LC720421</u>, and <u>LC699423</u>.

