## Supplemental Table S1 for "Evidence of the simultaneous replications of active viruses in specimens positive for multiple respiratory viruses"

Supplemental table S1. Primer/probe sequences used for the detection of respiratory viruses.

| Target (modification) | Primer/Probe names | Sequence (5' to 3') | Target | Length | Reference |
|---|---|---|---|---|---|
| hMPV | hMPV F | ARYTGCCRATCTTTGGBGTYATAG | fusion | 24 | (1) Modified |
| | hMPV R | TYTKACAATACCAYCCTTGRTCYTC | | 25 | |
| FAM-BHQ1 | hMPV Probe | MAARGCAGCYCCYTCTTGYTCMGRA | | 25 | |
| RSV | HRSV-F | ATGGCTCTTAGCAAAGTCAAGT | Matrix | 22 | (2), (3). |
| | HRSV-R | TGCACATCATAATTRGGAGTRTCA | | 24 | |
| FAM-**BHQ1**-Phosphate | HRSV A-P | ACACTCAACAAAGA**T**CAACTTCTRTCATCCAGCA | | 34 | |
| TEXsRED-**BHQ3**-Phosphate | HRSV B-P | ACATTAAATAAGGA**T**CAGCTGCTGTCATCCAGCA | | 34 | |
| HBoV | HBoV 2693f | GTCAACACAGAGCTTCCAATCC | NP-1 | 22 | (4) |
| | HBoV 2781r | TGAATTAGTACCATCTCTAGCAATGC | | 26 | |
| FAM-BHQ1 | HBoV 2724-TP | AGTGCCAGTAGAACCCACACCACCCT | | 26 | |
| Rhino | Pic-1 | TCCTCCGGCCCCTGAAT | 5' NTR | 17 | (5) |
| | Pic-3 | GAAACACGGACACCCAAAGTAGT | | 23 | |
| FAM-MGB | Pic-5-Probe | YGGCTAACCYWAACCC | | 16 | |
| HCoV (NL63) | HCoV-NL63_N-F | CTCTTTCTCAACCCAGGGCTG | Nucleocapsid | 21 | (6) |
| | HCoV-NL63_N-R | CGAGGACCAAAGCACTGAATAAC | | 23 | |
| FAM-BHQ1 | HCoV-NL63_N-Probe | ACCTCGTTGGAAGCGTGTTCCTACCA | | 26 | |
| HCoV (OC43) | HCoV-OC43_N-F | GGGTACTGGTACAGACACAACAG | Nucleocapsid | 23 | (6) |
| | HCoV-OC43_N-R | GGTGCCGTACTGGTCTTTAGC | | 21 | |
| Cy5-BHQ3 | HCoV- | CCGATGGCAACCAGCGTCAACTGCT | | 25 | |

| | | | | | |
|---|---|---|---|---|---|
| | OC43_N-Probe | | | | |
| HCoV (HKU1) | HCoV-HKU1_N-F | GTTGCTAATCACCAAGCTGACAC | Nucleocapsid | 23 | (7) |
| | HCoV-HKU1_N-R | CGTACCAGGCGGAAACCTAG | | 20 | |
| FAM-BHQ1 | HCoV-HKU1_N-Probe | CCCTCCGATGTTTCGTCAAGGGATCCT | | 27 | |
| HCoV (229E) | HCoV-229E-Foward | CAGTCAAATGGGCTGATGCA | Nucleoprotein | 20 | (8, 9) |
| | HCoV-229E-Reverse | AAAGGGCTATAAAGAGAATAAGGTATTCT | | 29 | |
| VIC-TAMRA | HCoV-229E-Probe | CCCTGACGACCACGTTGTGGTTCA | | 24 | |
| Adenovirus 2 (A, C, D and F)) | Ad2-F | CCAGGACGCCTCGGAGTA | Hexon | 18 | (10) |
| | Ad2-R | AAACTTGTTATTCAGGCTGAAGTACGT | | 27 | |
| FAM-BHQ1 | Ad2-probe | AGTTTGCCCGCGCCACCG | | 18 | |
| Adenovirus 4 (B and E) | Ad4-F | GGACAGGACGCTTCGGAGTA | Hexon | 20 | (10) |
| | Ad4-R | CTTGTTCCCCAGACTGAAGTAGGT | | 24 | |
| Cy5-BHQ3 | Ad4-probe | CAGTTCGCCCGYGCMACAG | | 19 | |
| HPIV1 | HPIV1_F | CCATCCTTTTTCTGCAATGTATCC | HN | 24 | (6) |
| | HPIV1_R | ATTGCAAACACTCTGATTAACATTGG | | 26 | |
| Cy5-BHQ3 | HPIV1_P | CGGTGGCTTAACAACTCCGCTCCAAGG | | 27 | |
| HPIV2 | HPIV2_F | GGACGCCTAAATATGGACCTCTC | HN | 23 | (6) |
| | HPIV2_R | GTGAGTGTAACACCAATGGGTCT | | 23 | |
| Cy5-BHQ3 | HPIV2_probe | CCCAGCTTTATCCCCTCAGCAACATCTCCC | | 30 | |
| HPIV3 | HPIV-3 F | ATGGACATGGCATAATGTGCTAT | HN | 23 | (6) |
| | HPIV-3 R | AATGCTYCCTGTGGGATTGAG | | 21 | |
| FAM-BHQ1 | HPIV-3 Probe | TCCCCATGGACATTCATTGTTTCCTGGTCT | | 30 | |
| HPIV4 | HPIV-4AB_F | CAAAYGATCCACAGCAAAGATTC | Nucleoprotein | 23 | (6) Modified |
| | HPIV-4AB_R | ATGTGGCCTGTAAGGAAAGCA | | 21 | |
| VIC-MGB | HPIV-4AB_P | GTATCATCATCTGCCAAAT | | 19 | |
| FluC | Flu C F | GACGACTACACACCAGACATCC | Matrix | 22 | (6) |

| | | | | | |
|---|---|---|---|---|---|
| | Flu C R | CTGAGACATTACTCCTGTATCTTTCAC | | 27 | |
| Cy5-BHQ3 | Flu C Probe | TTGCATCTCAACCAAGCTGTGATTGTTCCT | | 30 | |
| FluA | MP-39-67For | CCMAGGTCGAAACGTAYGTTCTCTCTATC | | 29 | |
| | MP-183-153Rev | TGACAGRATYGGTCTTGTCTTTAGCCAYTCCA | Matrix | 32 | (11) |
| FAM-MGB | MP-96-75ProbeAs | ATYTCGGCTTTGAGGGGGCCTG | | 22 | |
| FluB | NIID-TypeB TMPrimer-F1 | GGAGCAACCAATGCCAC | | 30 | |
| | NIID-TypeB TMPrimer-R1 | GTKTAGGCGGTCTTGACCAG | NS | 22 | (11) |
| FAM-MGB | NIID-TypeB Probe2 | ATAAACTTYGAAGCAGGAAT | | 23 | |

Red character shows labeled fluorescences (5’ to 3’)

8. Dare RK, Fry AM, Chittaganpitch M, Sawanpanyalert P, Olsen SJ, Erdman DD. Human coronavirus infections in rural Thailand: a comprehensive study using real-time reverse-transcription polymerase chain reaction assays. J Infect

Dis. 2007 Nov 1;196(9):1321-8.

9. Owusu M, Annan A, Corman VM, Larbi R, Anti P, Drexler JF, et al. Human coronaviruses associated with upper respiratory tract infections in three rural areas of Ghana. PloS one. 2014;9(7):e99782.

10. Wong S, Pabbaraju K, Pang XL, Lee BE, Fox JD. Detection of a broad range of human adenoviruses in respiratory tract samples using a sensitive multiplex real-time PCR assay. J Med Virol. 2008 May;80(5):856-65.

11. WHO. WHO information for the molecular detection of influenza viruses. 2021 [cited 2022 14 July]; Available from: https://cdn.who.int/media/docs/default-source/influenza/molecular-detention-of-influenza-viruses/protocols_influenza_virus_detection_feb_2021.pdf?sfvrsn=df7d268a_5
