## Supplemental Table S2 for "Evidence of the simultaneous replications of active viruses in specimens positive for multiple respiratory viruses"

Supplemental table S2. Virus replication pattern in multiple infections.

| Pattern | Positive viruses in specimen* | No. | Replicated virus | No. | Replicated virus | No. | Replicated virus | No. |
|---|---|---|---|---|---|---|---|---|
| 1 | ADV2/ADV4 | 2 | **ADV2/ADV4** | **1** | | | | |
| 2 | ADV2/ADV4/HRV | 1 | ADV4 | 1 | | | | |
| 3 | ADV2/ADV4/HMPV/HRV | 1 | HRV | 1 | | | | |
| 4 | ADV2/FluA/HRV | 1 | | | | | | |
| 5 | ADV2/FluA/HCoVHKU1/HMPV | 1 | | | | | | |
| 6 | ADV2/FluA/HMPV/HRV | 1 | | | | | | |
| 7 | ADV2/FluC/HBoV1 | 2 | | | | | | |
| 8 | ADV2/FluC | 2 | FluC | 1 | | | | |
| 9 | ADV2/HBoV1/HMPV | 1 | HMPV | 1 | | | | |
| 10 | ADV2/HBoV1/HMPV/HRV | 1 | **HMPV/HRV** | **1** | | | | |
| 11 | ADV2/HBoV1/HPIV3 | 1 | HPIV3 | 1 | | | | |
| 12 | ADV2/HBoV1/HPIV3/HRV | 1 | **ADV2/HRV** | **1** | | | | |
| 13 | ADV2/HBoV1 | 3 | ADV2 | 1 | | | | |
| 14 | ADV2/HBoV1/HRV | 3 | **ADV2/HBoV1** | **1** | **HBoV1/HRV** | **1** | | |
| 15 | ADV2/HBoV1/HRV/RSVA | 2 | **HRV/RSVA** | **1** | | | | |
| 16 | ADV2/HCoV229E/RSVB | 1 | | | | | | |
| 17 | ADV2/HCoVNL63/HMPV | 1 | HMPV | 1 | | | | |
| 18 | ADV2/HCoVOC43 | 4 | **ADV2/HCoVOC43** | **1** | HCoVOC43 | 1 | | |
| 19 | ADV2/HMPV/HRV | 1 | HRV | 1 | | | | |
| 20 | ADV2/HPIV3/RSVB | 1 | | | | | | |
| 21 | ADV2/HPIV3/HRV/RSVA | 1 | | | | | | |
| 22 | ADV2/HPIV4 | 5 | **ADV2/HPIV4** | **1** | HPIV4 | 1 | | |
| 23 | ADV2/HPIV4/RSVB | 1 | RSVB | 1 | | | | |
| 24 | ADV2/HRV | 3 | | | | | | |
| 25 | ADV2/HRV/RSVA | 1 | | | | | | |
| 26 | ADV4/HBoV1 | 1 | | | | | | |
| 27 | ADV4/HBoV1/HPIV3 | 1 | HPIV3 | 1 | | | | |
| 28 | ADV4/HBoV1/HPIV4 | 1 | ADV4 | 1 | | | | |
| 29 | ADV4/HBoV1/HPIV3/HRV/RSVB | 2 | HRV | 2 | | | | |
| 30 | ADV4/HBoV1/HCoVHKU1/HMPV/HPIV4/HRV | 1 | ADV4 | 1 | | | | |
| 31 | ADV4/HCoVOC43/HRV | 1 | | | | | | |
| 32 | ADV4/HMPV/HPIV3/RSVB | 1 | | | | | | |
| 33 | ADV4/HPIV1 | 1 | | | | | | |

| | | | | | | | | |
|---|---|---|---|---|---|---|---|---|
| 34 | ADV4/HPIV4 | 1 | | | | | | |
| 35 | FluA/HCoVHKU1 | 1 | FluA | 1 | | | | |
| 36 | FluA/HCoVNL63 | 1 | | | | | | |
| 37 | FluA/HPIV1/HRV | 1 | FluA | 1 | | | | |
| 38 | FluB/HRV | 1 | FluB | 1 | | | | |
| 39 | FluB/RSVB | 1 | FluB | 1 | | | | |
| 40 | FluC/HCoVOC43/RSVA | 1 | | | | | | |
| 41 | HBoV1/HCoV229E/RSVB | 1 | | | | | | |
| 42 | HBoV1/HCoVHKU1/RSVA | 1 | | | | | | |
| 43 | HBoV1/HCoVNL63 | 1 | HCoVNL63 | 1 | | | | |
| 44 | HBoV1/HCoVNL63/RSVA | 1 | **HBoV1/HCoVNL63** | **1** | | | | |
| 45 | HBoV1/HMPV | 3 | HMPV | 1 | | | | |
| 46 | HBoV1/HMPV/HPIV3/RSVA | 1 | HMPV | 1 | | | | |
| 47 | HBoV1/HPIV1 | 1 | | | | | | |
| 48 | HBoV1/HPIV3/HRV/RSVA | 1 | | | | | | |
| 49 | HBoV1/HPIV3/HRV | 3 | **HBoV/HPIV3/HRV** | **1** | HPIV3 | 1 | HRV | 1 |
| 50 | HBoV1/HRV | 6 | **HBoV1/HRV** | **3** | | | | |
| 51 | HBoV1/RSVA | 1 | RSVA | 1 | | | | |
| 52 | HCoV229E/HMPV | 2 | **HCoV229E/HMPV** | **1** | | | | |
| 53 | HCoV229E/HPIV3/RSVB | 1 | HPIV3 | 1 | | | | |
| 54 | HCoV229E/HRV/RSVA | 1 | HRV | 1 | | | | |
| 55 | HCoV229E/RSVB | 1 | | | | | | |
| 56 | HCoVHKU1/HCoVNL63/HCoVOC43 | 1 | | | | | | |
| 57 | HCoVHKU1/HCoVOC43 | 2 | HCoVOC43 | 2 | | | | |
| 58 | HCoVHKU1/HMPV | 1 | | | | | | |
| 59 | HCoVHKU1/HMPV/HRV | 1 | HCoVHKU1 | 1 | | | | |
| 60 | HCoVHKU1/HPIV3 | 1 | HPIV3 | 1 | | | | |
| 61 | HCoVHKU1/HPIV3/HPIV4/HRV/RSVA | 1 | | | | | | |
| 62 | HCoVNL63/HCoVOC43 | 1 | HCoVOC43 | 1 | | | | |
| 63 | HCoVNL63/HCoVOC43/RSVA | 3 | | | | | | |
| 64 | HCoVNL63/HRV | 1 | HCoVNL63 | 1 | | | | |
| 65 | HCoVOC43/HRV | 3 | **HCoVOC43/HBoV/HRV** | **1** | HCoVOC43 | 1 | | |
| 66 | HCoVOC43/HRV/RSVA | 1 | | | | | | |
| 67 | HCoVOC43/RSVA | 2 | RSVA | 1 | | | | |
| 68 | HMPV/HPIV1 | 1 | HPIV1 | 1 | | | | |
| 69 | HMPV/HRV | 3 | | | | | | |

| | | | | | |
|---|---|---|---|---|---|
| 70 | HMPV/HRV/RSVA | 1 | | | |
| 71 | HMPV/RSVA | 1 | | | |
| 72 | HMPV/RSVB | 1 | | | |
| 73 | HPIV1/HPIV4 | 1 | **HPIV1/HPIV4** | **1** | |
| 74 | HPIV1/HRV | 1 | HPIV1 | 1 | |
| 75 | HPIV2/HPIV3 | 1 | HPIV3 | 1 | |
| 76 | HPIV2/HPIV4 | 1 | HPIV4 | 1 | |
| 77 | HPIV2/RSVA | 1 | HPIV2 | 1 | |
| 78 | HPIV2/HCoVNL63/HPIV4/RSVA | 1 | | | |
| 79 | HPIV3/HPIV4/RSVA | 1 | | | |
| 80 | HPIV3/HPIV4/RSVB | 1 | HPIV3 | 1 | |
| 81 | HPIV3/HPIV4 | 2 | HPIV4 | 1 | |
| 82 | HPIV3/HRV | 1 | | | |
| 83 | HPIV4/HRV | 2 | HRV | 1 | |
| 84 | HPIV4/RSVA | 1 | | | |
| 85 | HPIV4/RSVB | 3 | HPIV4 | 1 | |
| 86 | HRV/RSVA | 2 | | | |
| | Total | 127 | | | 61 |

*, Virus names which was positive in specimen in diagnostic test .
